# GPSRdocker: A Docker-based Resource for Genomics, Proteomics and Systems biology

**DOI:** 10.1101/827766

**Authors:** Piyush Agrawal, Rajesh Kumar, Salman Sadullah Usmani, Anjali Dhall, Sumeet Patiyal, Neelam Sharma, Harpreet Kaur, Vinod Kumar, Dilraj Kaur, Shipra Jain, Akshara Pande, Sherry Bhalla, Gajendra P.S. Raghava

## Abstract

**Background:** In past number of web-based resources has been developed in the field of Bioinformatics. These resources are heavily used by scientific community to provide solution for challenges faced by experimental researchers particularly in the field of biomedical sciences. There are number of challenges in utilizing full potential of these services that includes internet speed, limits on computing power, and security of data. In order to enhance utilities of these web-based assets, we developed a docker-based container that integrates large number resources available in literature.

**Results:** This paper describes GPSRdocker a docker-based container developed for providing wide-range of computational tools in the field of bioinformatics particularly in genomics, proteomics and system biology. Majority of tools integrated in GPSRdocker are based on web services developed at Raghava’s group in last two decades. Broadly, these tools can be categorized in three categories; i) general scripts, ii) supporting software and iii) major standalone software. In order to facilitate students or developers working in the field of bioinformatics, we developed general scripts in Perl and Python. These general-purpose scripts serve as building block for any bioinformatics tools like computing features/descriptors of a protein. Supporting software packages includes SCIKIT, WEKA, SVM^light^, and PSI-BLAST; these software packages allow one to develop/implement bioinformatics software. Major Standalone software is core of this container which allows predicting function/class of biomolecules. These tools can be classified broadly in following categories; protein annotation, epitope-based vaccines, prediction of interaction and drug discovery.

**Conclusion:** A docker-based container has been developed which can be easily run on any operating system as well as it can be directly ported on cloud. Scripts can be run to build pipelines for addressing problems at system level like prediction of vaccine candidate for a pathogen. GPSRdocker including manual is available free for academic use from https://webs.iiitd.edu.in/gpsrdocker.

## 1. Introduction

Numerous software packages, libraries and web based services have been developed by bioinformaticians or computational biologist over the years. Only standalone software or libraries has been developed in pre-internet era, most of them were free for public use; mainly called domain software. Following archival sites have been created to maintain these software packages; EMBL File Serve [1], IUBIO archive [2], BioCatalog [3] and PDSB [4]. Our group developed, first scientific program ELISAeq [5] in 1990, for computing antigen/antibody concentration from ELISA data in GW-BASIC [6]. All these programs were standalone programs, developed for DOS/Windows using programming various languages like FORTRAN, PASCAL, C. These programs were distributed free for academic users via floppy, CD or via email-server. Though these programs were user-friendly, but one needs to have a hardware/software compatibility and knowledge of installation, in order to run these programs. In the era of Internet (1995 onwards), most of developers start to develop web based services. These webservers overcome limitations of standalone software packages where user only need to have a computer with browser and access to internet.

In last two decades, a wide range of web based services have been developed by scientific community particularly in the field of biology. Despite, numerous advantages of web based technology it has its own limitations. One of the major limitation of in the era of genomics is to predict function of all proteins or genes at genome level; transfer of huge data over internet is time consuming and costly. In addition, service provides cannot meet the computation requirement of user. Another concern is security of data, user do not wish to transfer confidential data over Internet. In order to provide service to community one need to use old technology of standalone software. There are number of challenges that include compatibilities of codes, dependencies, versions of libraries, compilers. This is nearly impossible for a user to install these software on their local machines. In order to overcome above limitations, number of projects have been initiated to develop customize operating system for free software in bioinformatics. These projects include BioLinux, VigyaanCD, DNAlinux, NEBC Bio-Linux, Vlinux. These operating systems are mainly flavors of Unix/Linux, like Ubuntu, Red Hat, Debian. A wide range of bioinformatics software packages have been integrated in these packages. Despite numerous advantages of customize operating systems, it consumes lot of computational resources.

In order to provide alternate to VMS, docker based and singularity based containers which are light weight and requires minimum resources became popular. Using these available container numbers of Bioinformatics pipelines have been developed (Eg. DNAp, NGSeasy, LncPipe etc.). In this manuscript, we have developed GPSRdocker (docker based container) that integrates bioinformatics software to perform various tasks. Most of the software in this package are unique and not integrated in any container or VMS so far. These software’s has been developed at Raghava’s group over the years and their webservers are available on the webpage (https://webs.iiitd.edu.in/gpsrdocker/). This software would be useful for researchers for developing various pipelines on their local machines.

## 2. Implementation

Docker provides a platform to perform operating system level virtualization or containerization. In brief, it provides a platform to develop, employ and run applications within a flexible and lightweight container. Containers are basically a software package, which are isolated from each other and pack their own configuration files, tools and libraries. All the containers can be interconnected for ease communication through well-defined channels and are actually run by a single operating system kernel. Containers are launched by running an image, which specify their precise contents. An image is basically the executable package constituting essentials needed to run a software i.e. code, libraries, configuration files and environment variables. GPSRdocker is a docker-based container that provides a resource on Genomics, Proteomics and System Biology. Concisely, GPSRdocker, is based on docker suite where customized container of all our webservers are available. User can run GPSRdocker on their machine using following steps

1. Install the docker into your system and create account at docker hub.
2. Make sure the docker is running before installing GPSRdocker.
3. Type following command install: **docker pull raghavagps/gpsrdocker**
4. Run docker on your machine using command: **docker run -i -t raghavagps/gpsrdocker**
5. Above step allow to work in docker image, now user can install software packages using command **“/gpsr/gpsr_install.pl”**. This will allow to users to install packages and provides instruction to run these software packages.

Complete manual on GPSRdocker is available from its web site https://webs.iiitd.edu.in/gpsrdocker/, user may read manual to install and run software packages.

### 2.1. General Scripts

In this section, we have described small programs developed at our group to generate features which can be used as building block to develop complex prediction modules. These programs are different from existing software libraries or modules like BioPERL, BioPython, as user should have programming skills in order to use these modules/subroutines. In GPSR 2.0 package we have developed small programs, which can be run by any person with minimal knowledge of programming skills. Following are important programs included in this package used to generate features for developing prediction models from protein and DNA/RNA sequences. These programs are developed in PERL and Python following the set standards. In order to run these codes, user needs to have Python 3.0 or above version installed in their system.

**Table 1:**
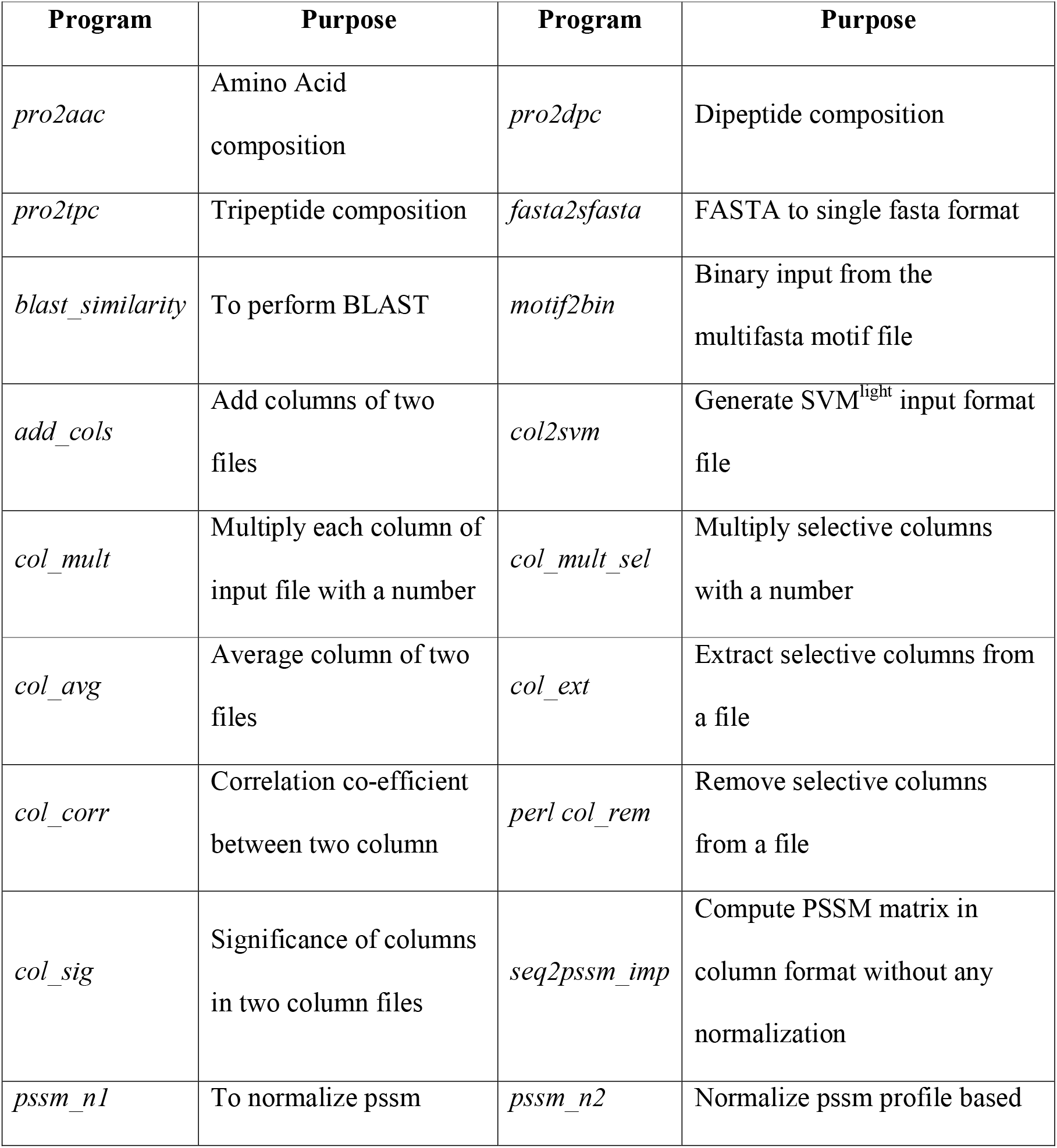
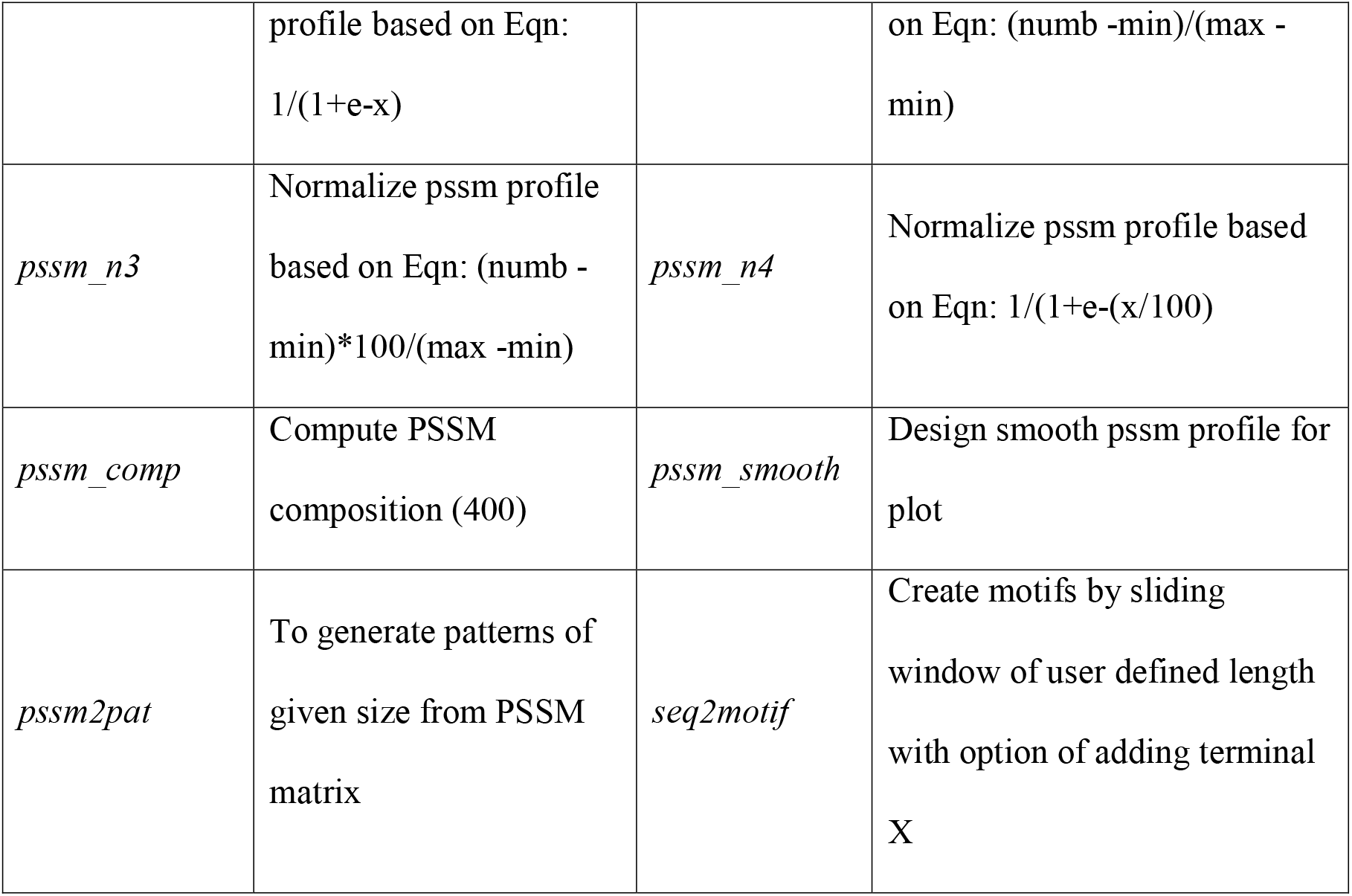
List of the scripts along with their purpose incorporated in the GPSRdocker.

| Program | Purpose | Program | Purpose |
| --- | --- | --- | --- |
| *pro2aac* | Amino Acid composition | *pro2dpc* | Dipeptide composition |
| *pro2tpc* | Tripeptide composition | *fasta2sfasta* | FASTA to single fasta format |
| *blast_similarity* | To perform BLAST | *motif2bin* | Binary input from the multifasta motif file |
| *add_cols* | Add columns of two files | *col2svm* | Generate SVM<sup>light</sup> input format file |
| *col_mult* | Multiply each column of input file with a number | *col_mult_sel* | Multiply selective columns with a number |
| *col_avg* | Average column of two files | *col_ext* | Extract selective columns from a file |
| *col_corr* | Correlation co-efficient between two column | *perl col_rem* | Remove selective columns from a file |
| *col_sig* | Significance of columns in two column files | *seq2pssm_imp* | Compute PSSM matrix in column format without any normalization |
| *pssm_n1* | To normalize pssm | *pssm_n2* | Normalize pssm profile based |
| | profile based on Eqn:<br>$1/(1+e^{-x})$ | | on Eqn: $(\text{numb} - \text{min})/(\text{max} - \text{min})$ |
| *pssm_n3* | Normalize pssm profile based on Eqn: $(\text{numb} - \text{min}) * 100 / (\text{max} - \text{min})$ | *pssm_n4* | Normalize pssm profile based on Eqn: $1/(1+e^{-(x/100)})$ |
| *pssm_comp* | Compute PSSM composition (400) | *pssm_smooth* | Design smooth pssm profile for plot |
| *pssm2pat* | To generate patterns of given size from PSSM matrix | *seq2motif* | Create motifs by sliding window of user defined length with option of adding terminal X |

### 2.2. Supporting Software

We utilized service of various software for developing and implementing our software. These supporting software include PSI-BLAST [7], CD-HIT [8], LPC [9], PSIPRED [10], machine learning packages like scikit-learn [11], SVM^light^ [12], SNNS [13], WEKA [14]. These software were used for data processing, developing dataset, performing alignment, removing redundancy among sequences, developing machine learning models and implementing them.

### 2.3. Standalone Packages

In this section, we are describing various standalone prediction packages developed in our group. For the ease of user we have classified this section into broad six categories such as 1) Protein Structure Prediction; 2) Functional annotation of proteins; 3) Vaccinomics; 4) Genomics: Genome annotation and application; 5) BioDrugs: Biomolecules based therapeutics; 6) Interactome: Biomolecular based therapeutics. We have categorized our prediction packages in these classes.

**Fig 1:**
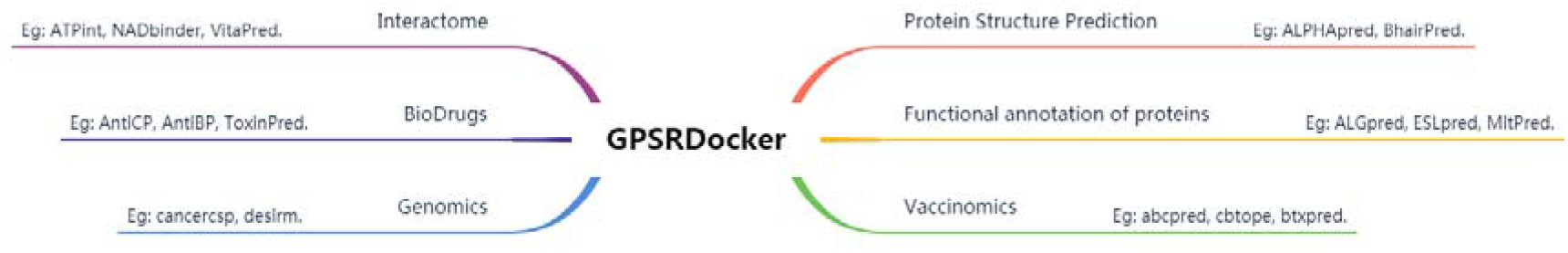
This figure shows various types of standalone packages in GPSRDocker.

#### 2.3.1. Protein Structure Prediction

Protein structure is traditionally determined using X-Ray crystallography, NMR spectroscopy and Cryo-electron microscopy. These methods are accurate but have certain limitations such as labor intensive, higher cost, etc. In order to bridge this gap, computational biologist have come up with *in silico* methods. *In silico* protein structure prediction is usually done by three methods (i) Homology Modelling; (ii) Threading based and (iii) ab initio method. In this section, we have described various methods developed by our group to predict 2D and 3D protein structure based on above principles. Described below are servers which can predict alpha turn, beta turns, gamma turns, phi-psi angle in protein, tertiary structure of proteins and peptides, etc.

**Table 2:**
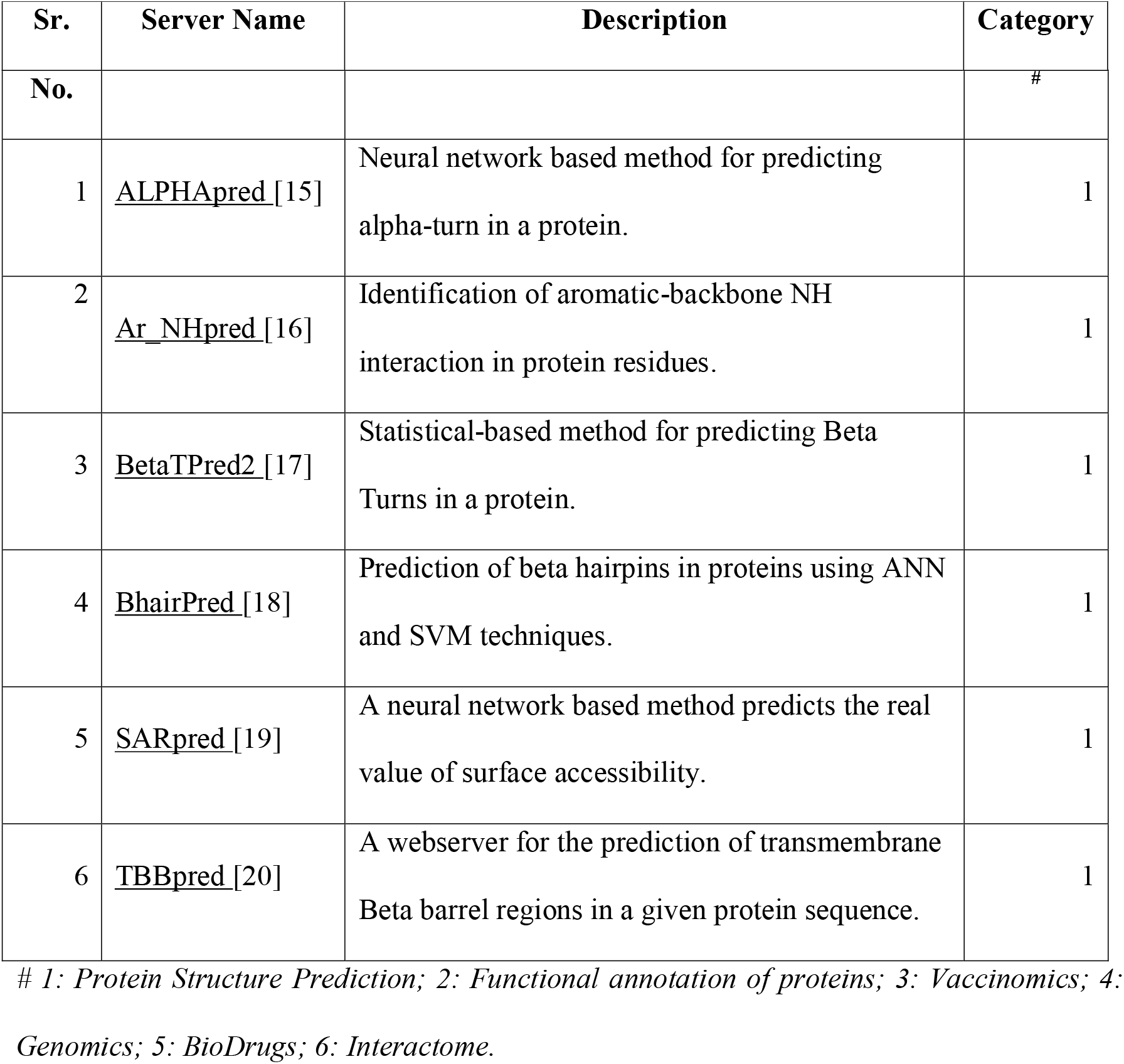
List of Protein Structure Prediction software incorporated in the GPSRdocker.

#### 2.3.2. Functional annotation of proteins

Proteins are key components in various biological processes. Protein interactions with other molecules in a biological system are responsible for signaling pathways, up regulation/down regulation of processes etc. Alteration or mutation in a protein sequence can lead to altered protein function, growth of various diseases, etc. Due to advancement in next generation sequencing techniques, large number of genome projects has been sequenced providing pool of protein sequences. However, functional annotation of these proteins is yet to be unfolded. Our group has developed number of tools to predict the function of proteins.

**Table 3:**
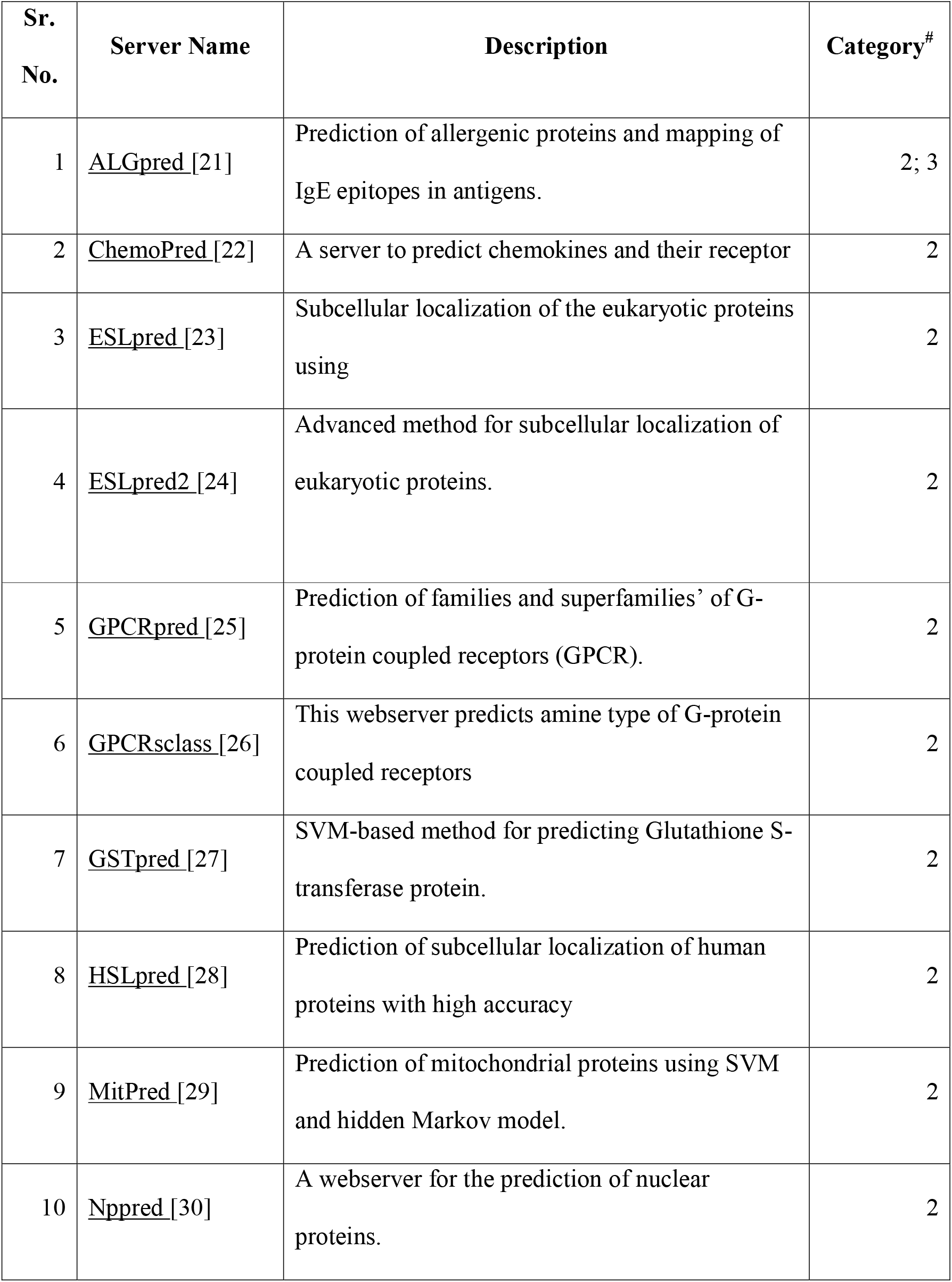
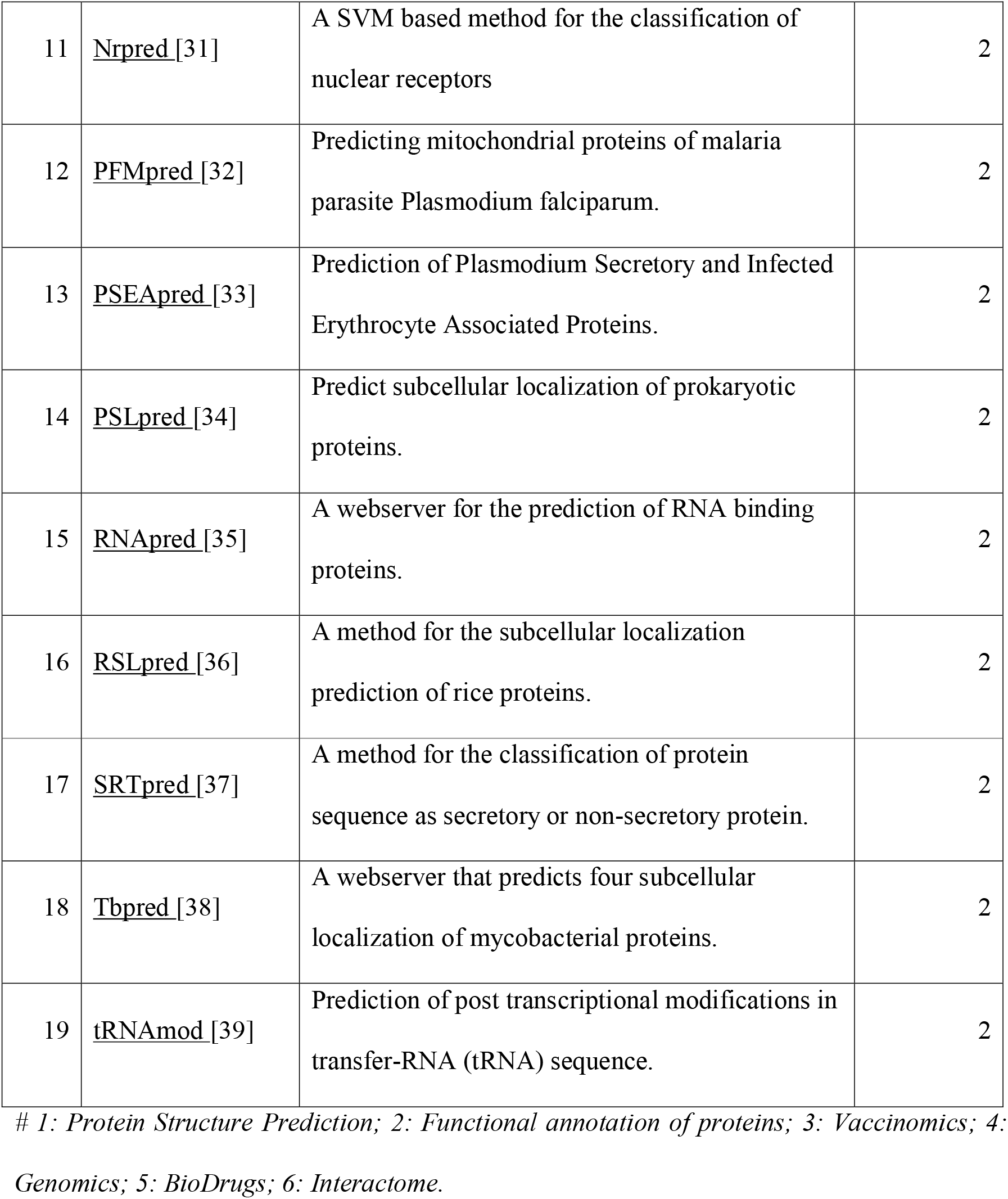
List of Protein Functional annotation software incorporated in the GPSRdocker.

#### 2.3.3. Vaccinomics

Vaccinomics combines immunogentics and immunogenomics with systems biology and immune profiling, to aid in understanding personalized or precision medicine. Vaccinomics is to comprehend biological immune system response towards vaccine-induced immunity of an individual. This paves the way for scientific community to design effective vaccines against hypervariable or resistant pathogens. Below are the tools developed by our group, which helps in development, administration and monitorization of potential vaccines.

**Table 4:** List of Vaccinomics software incorporated in the GPSRdocker.

| Sr. No. | Server Name | Description | Category<sup>#</sup> |
| --- | --- | --- | --- |
| 1 | abcpred [40] | Mapping of B-cell epitope(s) in an antigen sequence, using artificial neural network. | 3 |
| 2 | bcepred [40] | Prediction of linear B-cell epitopes, using Physico-chemical properties. | 3 |
| 3 | btxpred [41] | Prediction of bacterial toxins. | 3 |
| 4 | cbtope [42] | Conformational B-cell Epitope prediction. | 3 |
| 5 | ifnepitope [43] | Prediction and designing interferon-gamma inducing epitopes. | 3 |
| 6 | igpred [44] | Prediction of antibody specific B-cell epitope. | 3 |
| 7 | il10pred [45] | Prediction of Interleukin-10 inducing peptides. | 3 |
| 8 | il4pred [46] | In silico platform for designing and discovering of Interleukin-4 inducing peptides. | 3 |
| 9 | lbtope [47] | Prediction of linear B-cell epitopes. | 3 |
| 10 | pcleavage [48] | Identification of proteosomal cleavage sites in a protein sequence. | 3 |
| 11 | propred [49] | Prediction of MHC Class-II binding regions in an antigen sequence. | 3 |
| 12 | propred1 [50] | Prediction of promiscuous MHC Class-I binders. | 3 |
| 13 | tappred [51] | Prediction of binding affinity of peptides toward the TAP transporter. | 3 |
| 14 | vaxinpad [52] | Designing of peptide based vaccine adjuvant. | 3 |
| 15 | cancer_pred [53] | Prediction of the cancer lectins. | 3 |
| 16 | rnapin [54] | Prediction of Protein Interacting Nucleotides (PINs) in RNA sequences. | 3; 6 |

#### 2.3.4. Genomics: Genome annotation and application

With the advent of genomics era and next generation sequencing technologies, bioinformaticians have developed various tools for sequencing, assembling, structural and functional annotation of genomes. The sequencing data is increasing exponentially, and is available in public domain for analysis. Therefore, there is a need to develop tools which can effectively search, analyze and infer genomic information. In this direction, our group has developed many tools listed below:

**Table 5:** List of Genomics software incorporated in the GPSRdocker.

| Sr. No. | Server Name | Description | Category<sup>#</sup> |
| --- | --- | --- | --- |
| 1 | cancercsp [55] | Gene expression-based biomarkers for discriminating early and late stage of clear cell renal cancer. | 4, 3 |
| 2 | cancerspp [56] | Prediction and analysis of primary and metastatic tumor of SKCM using signature genes expression data. | 4, 3 |
| 3 | cancertsp [57] | Gene expression-based biomarkers for discriminating early and late stage of Papillary Thyroid Carcinoma (PTC). | 3 |
| 4 | desirm [58] | Designing of highly efficient siRNA with minimum mutation approach. | 4, 3 |

#### 2.3.5. BioDrugs: Biomolecules based therapeutics

BioDrugs also known as “bioactive drugs” are released in gastrointestinal tract by living orally administered recombinant microorganisms. These microorganisms are responsible for bioconversion or biosynthesis in the digestive environment. Bioactive drugs in model organisms like bacteria, yeasts etc. is a tedious and cost intensive experimental design. Chemoinformatics is the study of chemicals using various chemical databases, quantitative-structure activity relationship (QSAR), prediction of chemical properties or spectral. It plays a significant role in efficient drug discovery and development process. It has become an integral part of research in various fields like biochemistry, molecular biology, chemical genomics, bioinformatics etc. To facilitate this various bioinformatics tools have been developed which integrate experimentally validated data and used them to design new drugs. Our group has developed number of software’s to screen potential biodrugs in silico.

**Table 6:** List of Biodrugs software incorporated in the GPSRdocker.

| Sr. No. | Server Name | Description | Category<sup>#</sup> |
| --- | --- | --- | --- |
| 1 | AntiCP [59] | Prediction and design of anticancer peptides. | 5 |
| 2 | AHTpin [60] | Designing and virtual screening of antihypertensive peptides. | 5 |
| 3 | AntiBP [61] | Mapping of antibacterial peptides in a protein sequence. | 5 |
| 4 | AntiBP2 [62] | Advanced server for predicting antibacterial peptides with high precision. | 5 |
| 5 | CellPPD [63] | Computer-aided Designing of efficient cell penetrating peptides. | 5 |
| 6 | TumorHPD [64] | Server dedicated for designing tumor homing peptides. | 5 |
| 7 | HemoPI [65] | Prediction and virtual screening of hemolytic peptides. | 5 |
| 8 | ToxinPred [66] | An in silico method, which is developed to predict and design toxic/non-toxic peptides. | 5, 2, 3 |
| 9 | VICMPpred [67] | Prediction of Virulence factors, Information molecule, Cellular process and Metabolism molecule in the Bacterial proteins. | 5, 2 |
| 10 | NeuroPIpred [68] | Predict peptide as a neuropeptide or non neuropeptide. | 5 |
| 11 | AntiTbPred [69] | To predict peptides with bactericidal activity against Mycobacterium species. | 5 |

#### 2.3.6. Interactome: Biomolecular based therapeutics

Molecular interactions among biomolecules inside a cell are known as “Interactome”. Interactomics is the study of molecular interactions among proteins or small molecules and their consequences in the cell. These Bio-molecules could be proteins, nucleic acids, carbohydrates and lipid molecules. Interactomics could aid in identifying disease development and alteration in molecular mechanism of disease state. Below are few tools developed in our group to study interaction among biomolecules.

**Table 7:** List of Interactome software incorporated in the GPSRdocker.

| Sr. No. | Server Name | Description | Category<sup>#</sup> |
| --- | --- | --- | --- |
| 1 | ATPint [70] | Identification of ATP binding sites in ATP-binding proteins. | 6 |
| 2 | GlycoEP [71] | Prediction of C-, N- and O-glycosylation site in eukaryotic proteins. | 6,2 |
| 3 | GlycoPP [72] | Prediction of potential N-and O-glycosites in prokaryotic proteins. | 6,2 |
| 4 | GTPbinder [73] | Identification of GTP binding residue in protein sequences. | 6,2 |
| 5 | NADbinder [74] | Prediction of NAD binding proteins and their interacting residues. | 6,2 |
| 6 | <u>Pprint</u> [75] | ANN based method for identification of RNA-interacting residues in a protein. | 6 |
| 7 | <u>PreMieR</u> [76] | Identification of mannose interacting residues (MIRs) in protein sequences. | 6 |
| 8 | <u>VitaPred</u> [77] | Identification of different class of vitamin interacting residues in a protein. | 6 |
| 9 | SAMbinder [78] | A webserver for predict SAM interacting residue in a given protein sequence. | 6 |

*# 1: Protein Structure Prediction; 2: Functional annotation of proteins; 3: Vaccinomics; 4: Genomics; 5: BioDrugs; 6: Interactome.*

## 3. Applications of GPSRDocker

In this paper, we have launched GPSRdocker, which brings together various standalone versions of webservers developed by our group in various fields of bioinformatics. In this package we have tried to bring various open source software’s to serve scientific community in a user friendly manner. GPSRdocker provides number of applications in our scientific filed and these applications are discussed below in detail.

### (i) Development of Novel Therapeutic Pipelines

This is one of the biggest advantage of this docker container which provides an option to the user for developing new therapeutic pipelines. In the past decades, number of viruses and bacterial strains have been evolved which required immediate treatment to prevent their outbreak. User can implement different software present in the package for designing novel vaccine and drugs. User can utilize the genome of the new strains to identify the various epitopes (B-cell, T-cell and A-cell) which can be used as potential vaccine candidates. ZikaVR [79], EbolaVCR [80] and VacTarBac [81] are few examples of such type of pipelines.

### (ii) Cancer Risk stage prediction

The suite also comprises of packages like CancerCSP [55] which provides user to identify the potential biomarkers using gene expression data and predict the possible risk stage of a cancer patient. This will help user to start treatment faster.

### (iii) Annotating large amount of protein sequences

The image provides number of methods developed for predicting small molecule binding site in the protein sequence. The ligand includes ATP, GTP, NAD, FAD and SAM. These are important ligands used previously for designing drugs. User can annotate the protein function for its protein sequences by predicting the interacting site of these small ligands. Also, user can predict whether the protein sequence of their interest in nucleic acid binding (DNA & RNA), whether they are allergen proteins, etc. Packages like ESLpred2 [24], PSLpred [34] allows user to predict the subcellular localization of their proteins sequence.

### (iv) Designing novel therapeutic peptides

Number of packages have been incorporated into this docker image for designing various types of peptide therapeutics. User can design novel antimicrobial peptides such as antibacterial peptides, antifungal peptides, cell penetrating peptides, hemolytic peptides, antiangiogenic peptides, anticancer peptides, toxicity predicting peptides, chemically modified cell penetrating peptides, chemically modified antimicrobial peptide, etc.

### (v) Easy to use on large dataset

One of the main problems with the web-based services is that they are not able to process after a certain limit and size of the dataset. GPSRdocker allows user to implement and use the same service on the large dataset without any issue of file size and space. User needs to check the space availability of its local machine and thereafter can use the package the way he wants too.

### (vi) Data security

Data security is one of the majors concerned nowadays. User can use the standalone service for securing its data and without providing any personal details. Also user can store any amount of data in the image without any loss of information by following the standard protocols while working in the docker container. User can also keep the data safe by saving the data in another image.

### (vii) No internet requirement

internet availability is prerequisite for accessing the web based service. However, in case of docker standalone package, there is no requirement of the internet once the image is pulled on the local machine. User can work anywhere in the container without the internet presence.

### (viii) Comprehensive resource of software

GPSRdocker provides a comprehensive resource of software related to different field of science such as immunoinformatic, protein structure and function annotation, cheminformatics, bio drugs, vaccine designing, genomics, etc. To the best of author’s knowledge there is no such platform developed previously which comprises of more than 60 standalone version of the software. Therefore, this package will be very useful to the researchers both working in the wet lab as well as in dry lab. The package could be useful to various pharmaceutical companies as well as to the students who are starting their career in the area of bioinformatics.

## 4. Conclusion

GPSRdocker is a user friendly docker based container developed by our group which can be used to run standalone versions of various web servers. At present, GPSRdocker contains around 65 standalone software of the webservers developed by our group which are highly cited in the literature. Each server included in this container is used to address various questions in the field of computational biology. Aim of developing GPSRdocker is to integrate various freely available resources on a platform which is compatible with all type of operating systems. With the rapid advancement in the field of bioinformatics, there is a need to implement cloud based technologies such as Docker to make resources easily accessible to the users. The only limitation of this work is that it includes software developed specifically in our group only. However, there are various other useful bioinformatics containers available in market. We are working to include all the possible general bioinformatics modules as well as other new bioinformatics webservers in GPSRdocker version2.

## Acknowledgements

Authors are thankful to funding agencies Department of Biotechnology (DBT) and Department of Science and Technology (DST-INSPIRE) and Council of Scientific and Industrial Research (CSIR), Govt. of India and Indraprastha Institute for Information Technology for financial support and fellowships.

## Author contribution

PA, SP, AD, NS, AP, RK, VK and DK developed the python codes. PA, SSU, RK, AD, SP, NS, HK, VK, DK, SJ, and AP developed the standalone versions of the software. PA, SSU, AD, SP, NS, DK, SJ and VK developed the tables and figure. RK and VK developed the website and manual was written by PA, SSU and GPSR. PA, SJ, HK, and GPSR wrote the manuscript. GPSR conceived the idea and coordinated the project. All authors read and approved the final paper.

